# Multiple longitudinal tracts in the cephalopod arm sensorimotor system

**DOI:** 10.1101/2025.11.20.689621

**Authors:** Cassady S. Olson, Clifton W. Ragsdale

**Author notes:** **Corresponding author:** Cassady S. Olson.

## Abstract

Octopuses have an incredibly rich behavioral repertoire, exhibiting complex motor acts that require the coordination of eight highly flexible arms, each with hundreds of suckers. These movements are controlled by an axial nerve cord (ANC), equivalent to the spinal cord, situated in the center of the arm musculature. The ANC has a cell body layer which forms a U-shape around its neuropil and is capped aborally, or opposite the sucker, by the cerebrobrachial tract (CBT), a massive fiber bundle known to interconnect the arms and the brain. In vertebrate spinal cords, in addition to the major fiber tracts that interconnect the brain and spinal cord, there are spinospinal connectives that coordinate complex motor behaviors across the appendages. Here, we asked with tract-tracing and immunohistochemistry, whether an octopus arm’s ANC might also have intrinsic longitudinal connections for coordinated arm and sucker movements. We found that the ANC neuropil is enriched in longitudinal fibers. These fibers form distinct tracts, two within the oral (sucker-side) neuropil and two in the aboral (brachial-side) neuropil. In addition, CBT itself demonstrates four major subtracts, and DiI labeling and dextran tracing suggests that (1) the CBT also carries arm-intrinsic longitudinal connections and (2) the CBT and the neuropil tracts can be subcategorized into those that primarily connect with the sucker and those that serve the arm musculature. We also examined the organization of fiber-tracts in the ANC of the arms and tentacles of two species of squid, establishing that an aboral, extra-neuropil tract is a shared feature across all cephalopod species studied. In addition, the squids also had an oral longitudinal tract, though its positioning and size varied with species and appendage. In sum, these findings describe the neural substrate for coordinating motor behaviors across the length of a cephalopod appendage.

## Introduction

Octopuses possess a motor control challenge of enormous complexity (Hochner et al., 2023; Olson and Ragsdale, 2023). Each of the hydrostatic arms move in near infinite degrees of freedom, and each arm is lined with hundreds of suckers, which also move independently (Kier and Stella, 2007; Kennedy et al., 2020; Röckner et al., 2023). Critical behaviors from locomotion to feeding require the octopus to coordinate motor activity along the length of an arm and between arms (Gutfreund et al., 1998; Sumbre et al., 2006; Kennedy et al., 2020; Bidel et al., 2022; Bennice et al., 2025). Furthermore, many of these behaviors require the coordination of sucker movements and the collaboration of motor activity between the suckers and the arm (Grasso, 2008; Kennedy et al., 2020; Buresch et al., 2022, 2024; Bennice et al., 2025).

Neural control for arm and sucker movements arises from the extensive nervous system embedded in the arm itself and from the intermediate motor centers in the octopus brain (Fig. 1a; Graziadei, 1971; Young, 1971; Hochner et al., 2023; Olson and Ragsdale, 2023). Each arm houses an axial nerve cord, equivalent to the vertebrate spinal cord, situated in the center of the brachial musculature (Fig. 1a-d; Graziadei, 1971; Neacsu and Crook, 2024; Olson et al., 2025). The ANC contains a cell body layer (CBL) that wraps around its neuropil (NP) (Fig. 1b, c). The CBL comprises repeated segments down the long axis of the arm, and, orthogonal to this architecture, the CBL and NP can be split into an oral, sucker territory and an aboral brachial territory (Rowell, 1963; Olson et al., 2025). The oral sucker territory of the ANC forms local enlargements for each sucker (Rossi and Graziadei, 1954; Olson et al., 2025). While ANC segments in the CBL and the division for brachial and sucker territories provide the structure for parsing the arm into smaller units for local control, they do not account for how movements of the suckers and the arm itself over a distance are coordinated.

**Figure 1:**
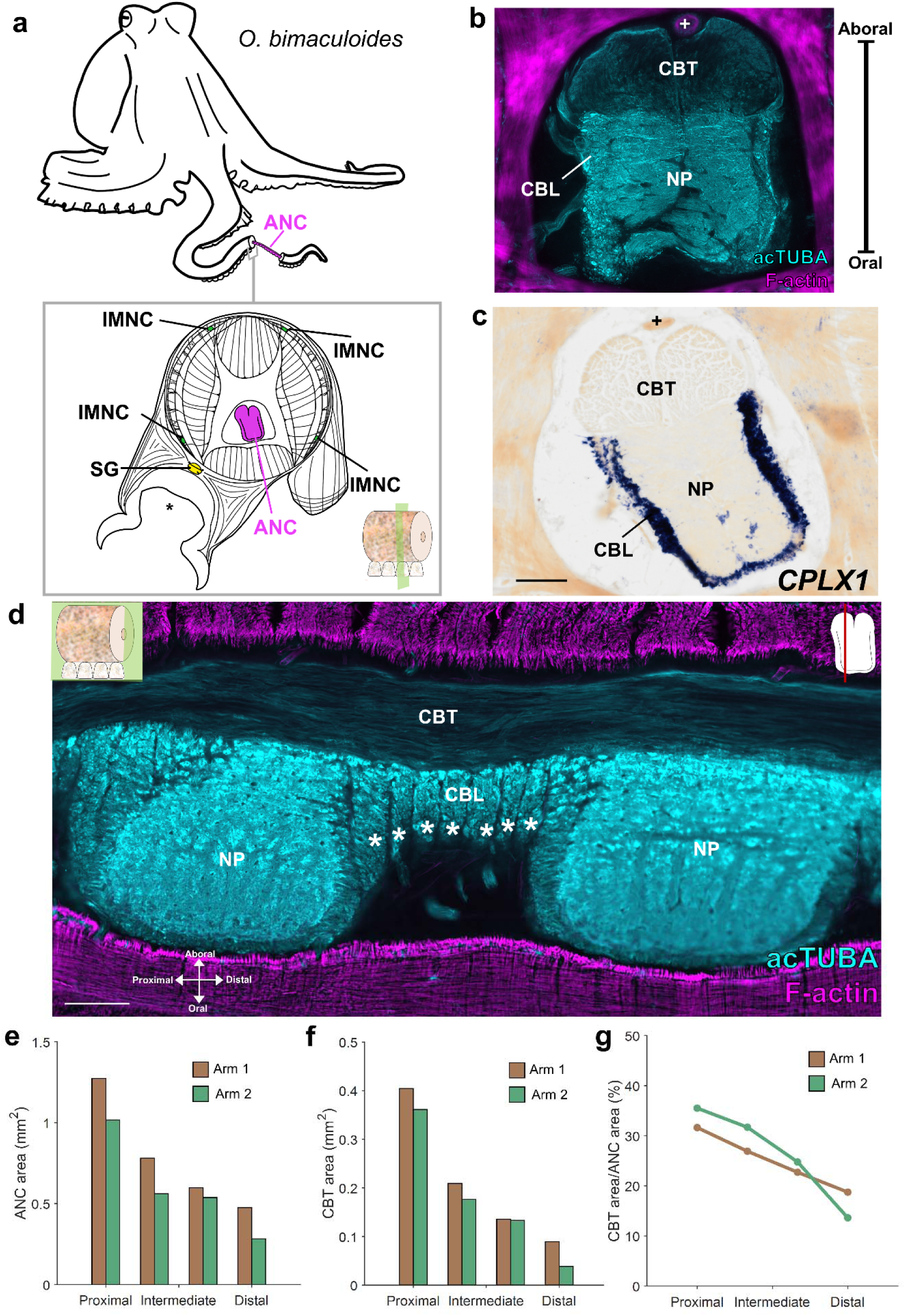
Cerebrobrachial tract composition and proximal-distal variation. **a,** Cartoon of *Octopus bimaculoides* and diagram of a transverse cross section of octopus arm with the major components of the arm nervous system highlighted: magenta, axial nerve cord (ANC); green, intramuscular nerve cord (IMNC); yellow, sucker ganglion (SG). Key indicates transverse plane of section. * denotes acetabulum of the sucker. **b,** Transverse section of the ANC stained for acetylated α-tubulin (acTUBA, cyan) and F-actin (magenta). The cerebrobrachial tract (CBT) is aboral (away from the sucker) to the cell body layer (CBL) and neuropil (NP) and is composed of acTUBA-positive fibers. The brachial artery, denoted by +, is situated in between the two CBT trunks. **c,** In situ hybridization (ISH) for complexin 1 (*CPLX1)* gene expression in a transverse section of the ANC labels neuronal cell bodies. Notably, the CBT is devoid of labeling. + denotes brachial artery. Scale bar: 250 μm. **d,** Longitudinal section of the ANC stained for acTUBA (cyan) and F-actin (magenta). CBT fibers are oriented longitudinally. CBL segments, denoted by white asterisks, are readily apparent in this preparation. Key indicates plane of section. Scale bar: 250 μm. **e, f,** Cross sectional area of the ANC (**e**) and the CBT (**f**) decreases from proximal to distal. The total cross-sectional area of the ANC = CBL area + NP area + CBT area. **g**, The proportion of total ANC area occupied by the CBT decreases from proximal to distal. acTUBA, acetylated α-tubulin; ANC, axial nerve cord; CBL, cell body layer; CBT, cerebrobrachial tract; IMNC, intramuscular nerve cord; NP, neuropil; SG, sucker ganglion.

In vertebrate nervous systems, major fiber tracts serve to interconnect the brain and spinal cord, allowing for the control and coordination of complex motor acts. These tracts can be divided into ascending sensory tracts, transmitting sensory input from the periphery, and descending motor tracts, relaying motor commands to spinal cord motor neurons. Analogously, the octopus ANC features a cerebrobrachial tract (CBT) situated aboral to the CBL and NP (Graziadei, 1971; Zullo et al., 2019; Neacsu and Crook, 2024). The CBT comprises two massive trunks of longitudinally running fibers that interconnect the arms and the brain (Graziadei, 1971; Zullo et al., 2019). As the ANC ascends towards the brain, the CBT coalesces into a single trunk (Graziadei, 1971). Classic studies have noted a parcellation of CBT fibers based on size, though their relative functions are unknown (Graziadei, 1971). Recent electrophysiological experiments posit that the CBT transmits motor signals *en passant,* allowing for an interplay between signals from the brain and ANC circuitry (Zullo et al., 2019). These studies are, however, few in number, and there is little experimental evidence for how longitudinally running tracts in the CBT are organized.

The vertebrate spinal cord also contains spinospinal connections, which are particularly prominent, and in fact quite extensive, in four-legged carnivores (Sherrington and Laslett, 1903). These spinospinal fibers travel in spinal cord white matter, thereby invading the major ascending and descending fascicles. Similarly, in the octopus system, classic studies suggest shorter interconnecting systems within the NP of the ANC, which most certainly assist in behaviors coordinating suckers and arm movements along a single arm (Graziadei, 1971). The relative positioning and trajectories of these inter-NP tracts are not known.

Here, we employ tract-tracing methods and immunohistochemistry to investigate the organization of CBT in *Octopus bimaculoides*. We further interrogate the ANC NP for additional longitudinal fiber tracts. Finally, we examined the organization of fiber-tracts in the ANC of the arms and tentacles of two species of squid to ask what the shared features of coleoid appendage fiber-tracts are. Our findings establish the presence of multiple longitudinal fiber tracts, in both the CBT and the NP of the ANC, that are strong candidates to mediate longer range movement coordination.

## Materials and Methods

### Animals

Aquatic Research Consultants, a field-collection venture operated by Dr. Chuck Winkler, San Pedro, CA, supplied wild caught adult *Octopus bimaculoides* (*Obi,* n = 17) of both sexes and at least six months of age. Octopuses were individually housed in 20-gallon saltwater tanks equipped with an UV light, carbon particle filter, and an aquarium bubbler. Fiddler crabs, bivalves or frozen shrimp were fed to the octopuses daily. The artificial seawater (ASW) was prepared by adding pharmaceutically pure sea salt (33g/liter; Tropic Marin “classic”, Wartenberg, Germany) to deionized water. Wild caught adult *Doryteuthis pealeii* (*Dpe,* n = 3) and lab cultured adult *Euprymna berryi* (*Ebe,* n = 3) of both sexes were obtained at the Marine Biological Laboratory, Woods Hole, MA.

### Tissue collection

Octopuses were deeply anesthetized in 4% ethanol/ASW (EtOH/ASW, n = 9) and trans-orbitally perfused with 4% paraformaldehyde/phosphate-buffered saline solution (PFA/PBS, pH 7.4) delivered via a peristaltic pump (Cole Parmer Masterflex) through a 21½ gauge needle (Becton Dickinson). The left and right white bodies, a hematopoietic tissue, were each targeted multiple times throughout the procedure. Arms were dissected and stored in 4% PFA/PBS overnight at 4°C. Specimens were then either cut into 2-4 cm pieces for immediate processing or stored in diethyl pyrocarbonate (DEPC) treated-PBS at 4°C.

Squid were kept in circulating, filtered containment tanks for several days before being deeply anesthetized in 7.5% MgCl_2_/ASW and dissected (*Dpe*, n = 3; *Ebe*, n = 3). Arm crowns were immersion-fixed in 4% PFA/PBS. Arms and tentacles were dissected and cut into 2-4 cm pieces for further processing.

### Slide preparation

Octopus and squid tissue blocks were placed in 30% sucrose/4%PFA/PBS at 4°C until saturated (three to five days), rinsed with 30% sucrose/PBS, and infiltrated with 10% gelatin/30% sucrose/PBS for 1 hour at 50°C. Tissue was embedded in 10% gelatin/30% sucrose/PBS and post-fixed in 30% sucrose/4%PFA/PBS before storage in −80°C. Serial arm sections were cut at 28-50-µm thickness on a freezing microtome (Leica SM2000R) and collected in DEPC-PBS. Sections were mounted and dried on charged, hydrophilic glass slides (TruBond380, Newcomer Supply, Middleton, WI), then stored at −80°C until further processing.

### In situ hybridization

To generate antisense riboprobes for *CPLX1,* plasmids were linearized through SpeI (New England Biolabs, Cat#: 50811989) restriction enzyme digestion to make template. Following phenol-chloroform extraction of the template, antisense digoxigenin (DIG)-labeled riboprobes (Sigma-Aldrich, Cat#: 11277073910) were transcribed with T7 RNA polymerase (New England Biolabs). After transcription, residual template was digested with RNase-free DNase I (Sigma-Aldrich) at 37°C for 15 minutes. Riboprobes were ethanol-precipitated and stored in 100μl of DEPC-H2O at −20°C until use. The cloning for *CPLX1* was previously reported (Olson et al., 2025)

Slides of sectioned octopus tissue were equilibrated to room temperature, then post-fixed in mailers for 15 minutes in 4% PFA/PBS, washed 3x 15 minutes (3 times for 15 minutes each) in DEPC-PBS and incubated at 37°C for 15 minutes in proteinase K solution (Sigma-Aldrich; 1.45 μg proteinase K per milliliter of 100 mM Tris-HCl [pH 8.0], 50 mM EDTA [pH 8.0]). Following a 15-minute post-fix in 4% PFA/PBS, slides were washed 3x 15 minutes in DEPC-PBS and acclimated to 72°C for one hour in hybridization solution (50% formamide, 5x SSC, 1% SDS, 200 μg/ml heparin, 500 μg /ml yeast RNA). Slides were transferred to mailers with 1-2 mg antisense riboprobe in 15 mL hybridization solution and incubated overnight at 72°C. The next day, slides were treated 3x 45 minutes in preheated Solution X (50% formamide, 5x SSC, 1% SDS) at 72°C. Slides were washed 3x 15 minutes in room temperature TBST (Tris-buffered saline with 1% Tween 20) and blocked at room temperature for 1 hour in 10% DIG buffer (Roche) in TBST.

Anti-DIG Fab fragments conjugated to alkaline phosphatase (Sigma-Aldrich, Cat#: 11093274910; RRID: AB_514497) were pre-absorbed with octopus embryo powder in 1% DIG buffer in TBST for at least one hour. Slides were then incubated on a rocker overnight at 4°C in preabsorbed antibody diluted to a final concentration of 1:5000 in 10% DIG buffer in TBST.

The next day, slides were washed once, then for 3x 15 minutes, then for 2x 1 hour in TBST. Slides were placed in freshly prepared NTMT (100 mM Tris-HCl [pH 9.5], 100 mM NaCl, 50 mM MgCl2, 1% Tween 20) for 10 minutes. For the color reaction, slides were incubated in 0.35 mg/mL nitro blue tetrazolium (NBT, stock: 100mg/mL in 70% dimethyl formamide/30% DEPC-H20, Gold Biotechnology, St. Louis, MO) and 0.175 mg/mL 5-bromo-4-chloro-3-indolyl phosphate (BCIP, stock: 50mg/mL in 100% dimethyl formamide, Gold Biotechnology, St. Louis, MO) in NTMT. Color development proceeded at room temperature and was monitored for a maximum of 5 days. When the reaction appeared complete, slides were washed in TBST overnight and dehydrated through a series of ethanol washes into Histoclear, and coverslipped with Eukitt (Sigma-Aldrich).

### Section and wholemount immunohistochemistry

Neuronal processes in the octopus arm and the squid arm and tentacle were labeled with mouse monoclonal antibody 6–11B-1 (1:500 dilution of ascites fluid; Sigma-Aldrich, Cat#: T6793; RRID: AB_477585) and mouse monoclonal antibody SMI-31 (1:500 dilution; BioLegend, Cat# 801601; RRID: AB_2564641). Clone 6–11B-1 was isolated following immunization with sea urchin sperm flagella protein preparations (Piperno and Fuller, 1985). It recognizes an acetylated α-tubulin (acTUBA) epitope found broadly but not universally across microtubules and has been extensively employed to identify axon tracts in vertebrate and invertebrate nervous systems (Chitnis and Kuwada, 1990; LeDizet and Piperno, 1991; Shigeno and Yamamoto, 2002; Baratte and Bonnaud, 2009; Olson et al., 2025; 2025a). Clone SMI-31 reacts with a phosphorylated epitope of neurofilament in mammals and neurofilament 220 in squid (Sternberger and Sternberger, 1983; Grant et al., 1995; Grant and Pant, 2016), and it has been employed successfully in octopus arm tissue (Olson et al., 2025).The secondary Alexa Fluor® 488 AffiniPure Goat Anti-Mouse IgG (Cat#: 115-545-003; RRID: AB_2338840) and Cy™3 AffiniPure Donkey Anti-Mouse IgG (Cat#: 715-165-151; RRID: AB_2315777) were purchased from Jackson ImmunoResearch (West Grove, PA) and utilized at 1:500 dilutions.

For section immunohistochemistry, slides were rinsed 3x 10 minutes in DEPC-PBS containing 1% Tween 20 (PBST) and incubated for 30 minutes in a proteinase K solution (Sigma-Aldrich, Cat#: 03115828001; 19.4 μg proteinase K per milliliter of PBST). Slides were post-fixed with 4% PFA/PBS for 15 minutes, washed 3x 30 minutes in PBST, blocked in 10% goat serum/PBST (Fisher Scientific, Cat#: 16210072) for 1 hour and incubated in primary antibody diluted in 1% goat serum/PBST for four days at 4°C. After 3x 30 minutes PBST washes, sections were incubated for two days at 4°C with secondary antibody along with 0.01 mg/ml 4′-6-diamidino-2-phenylindole (DAPI; Sigma-Aldrich, Cat#: D9542) to label cell nuclei fluorescently and either 0.2 µl/ml phalloidin-iFluor 594 (Abcam, Cat#: ab176757) or phalloidin-iFluor 488 (Abcam, Cat#: ab176753) to label filamentous actin (F-actin). Slides were washed 3x 5 minutes with PBST and cover slipped with Fluoromount G (Southern Biotech, Birmingham, AL).

For whole mount immunohistochemistry, the following octopus arm blocks were prepared: 0.5 cm – 1 cm transverse slices, or longitudinally bisected slices, or horizontally bisected slices. Slices were washed 3x15 minutes in PBST, dehydrated in a graded methanol series (25, 50, 75% in PBST, each for 10 minutes), rinsed twice in 100% methanol for 10 minutes, and stored overnight at −20°C. The next day, slices were rehydrated in a graded methanol series (75%, 50%, 25% in PBST, each for 5 minutes) and rinsed twice in PBST for 5 minutes. Then, the tissue was incubated for 60 minutes at 37°C in a proteinase K solution (19.4 μg proteinase K per milliliter of PBST), post fixed for 15 minutes in 4% PFA/PBS, washed 3x 15 minutes and 1x 1 hour in PBST, and blocked for an hour in 10% goat serum/PBST. Slices were incubated in primary antibody for 7 days at 37°C. Following primary incubation, tissue was rinsed 1x, then 3x 15 minutes, then 5x 1 hour and then overnight in PBST. Tissue was transferred to secondary antibody for seven days at 4°C. The tissue was rinsed with PBST 1x quickly, 3x 15 minutes, 2x 1 hour, washed with DEPC-PBS 3x 15 minutes, and post fixed for four days at 4°C. Slices were then rinsed 3x 15 minutes in PBS and cleared in a modified version of FRUIT (35%, 40%, 60%, 80% and 100% FRUIT, each for 24 hours; Hou et al., 2015). Slices were stored in 100% FRUIT at 4°C until imaged

### DiI labeling

DiI crystals (Biotium, Cat#: 60016) were placed at various targets in the 0.5-1cm transverse, horizontally and longitudinally bisected fixed arm blocks. Slices were incubated in 1% PFA/PBS in the dark for 7 days to 6 months. Following DEPC-PBS washes for 3x 15 minutes, explants were cleared in a modified version of FRUIT (Hou et al., 2015).

### Tracing

Adult *O. bimaculoides* (n = 10) were anesthetized in 2% EtOH/ASW, and the distal portion of a single arm was amputated with a razor blade. The amputated arm was cut with a razor blade into 0.5 cm-1 cm transverse slices. Slices were placed into 221 media with NuSerum (36% Leibovitz L15-media, 36% filtered ASW, 18% deionized water, 1% pen-strep, 10% NuSerum) prior to injection. Each slice (n = 103) was injected with CF® 488A Dye Dextran (Biotium, Cat#: 80110; 2% solution, 10kD) using a 25- or 32-gauge Hamilton syringe (Hamilton, model#: 7000.5), rinsed in filtered ASW, and incubated at room temperature in 221 media with NuSerum. After three hours, the slices were rinsed with filtered ASW and fixed in 4% PFA/PBS. Slices were gelatin embedded as described above, sectioned at 50µm, and counterstained with phalloidin-iFluor 594 (0.2 µl/ml in 1%PBST) and DAPI (0.01 mg/ml in 1% PBST) before imaging.

### Imaging

The immunolabeled and tracing tissue was studied with a Zeiss Axioskop 50 upright microscope and a Leica MZ FLIII stereomicroscope, both outfitted with the Zeiss AxioCam digital camera and AxioVision 4.5 software system. Selected sections were also studied on a Leica SP5 Tandem Scanner Spectral 2-photon confocal microscope (Leica Microsystems, Inc., Buffalo Grove, IL), a 3i Marianas Super-resolution through Optical Reassignment (SoRa) Yokogawa-type spinning disk confocal microscope (Intelligent Imaging Innovations, Denver, CO) with dual Prime 95B Illuminated Scientific CMOS (Teledyne Photometrics Vision Solutions, Tucson, AZ) run by Slidebook software (Intelligent Imaging Innovations, Denver, CO), or scanned with an Olympus VS200 Research Slide Scanner (Olympus/Evident, Center Valley, PA) with a Hamamatsu ORca-Fusion camera (Hamamatsu Photonics, Skokie, IL). Whole mounts were imaged on a LaVision BioTec UltraMicroscope II (Miltenyi Biotec, Bergish Gladbach, Germany) run by ImSpector Pro v. 7_124 software (LaVision BioTec, Bielefeld, Germany). Collected images were corrected for contrast and brightness and false colored in FIJI (version 2.1.0/1.53c; National Institutes of Health (NIH)). Diagrams were drawn using GIMP v. 3.0.41.0.

### Cerebrobrachial tract (CBT) proximal-distal analysis

To study proximal-distal variation in the CBT, four wholemount slices spanning the proximal-distal axis of one arm from two different animals (Arms 1L and 3R) were immunostained for acTUBA. Transverse confocal stacks were imaged at 10x to capture the whole slice and at 20x to capture the CBT. Maximum projections of the CBT were used to determine CBT subdivisions. Maximum projections of the whole slice were used to determine area.

#### CBT Subdivisions

Across all slices, discrete bundles of fibers within the CBT were outlined. Bundles that were consistent along the proximal-distal axis and between arms were demarcated as the main subdivisions.

#### CBT Area

The areas of the CBT and the ANC were calculated using the polygon tool in FIJI. To ensure the measurement was accurate, adhering the contours of the amorphous octopus anatomy, tracing dots were placed as close to each other as possible. The area for the CBT was then compared to the area of the ANC (CBT/ANC).

## Results

### Organization of the cerebrobrachial tract

Within the ANC of *O. bimaculoides*, the CBT comprises two trunks of longitudinally arranged nerve fascicles situated at the most aboral portion of the ANC (Fig. 1b, d). The brachial artery marks the midline between the two trunks (Fig. 1b; Graziadei, 1971; Olson et al., 2025). As demonstrated by in situ hybridization with the pan-neuronal marker *CPLX1* and with immunostaining for acTUBA, the CBT is devoid of neuronal cell bodies (Fig. 1b-d). The arm tapers down the long axis, with the suckers, brachial musculature, and axial nerve cord all decreasing in size (Fig. 1e; Olson et al., 2025). The area of the CBT likewise decreases from proximal to distal (Fig. 1f). Moreover, the proportion of the total ANC area occupied by the CBT decreases from proximal to distal (Fig. 1g). This decrease in fiber tract size is reminiscent of vertebrate spinal cord tracts, which are largest closest to the brain (Parent and Carpenter, 1996), and presumably reflects the decrease in fibers projecting to and from the brain as one moves away from the brain.

Using transverse whole mounts immunostained for acetylated α-tubulin (acTUBA), we identified subdivisions in the CBT trunks. Four distinct bundles could be recognized along the proximal-distal axis in the two arms studied (α, β, γ, and δ tracts; Fig. 2). These bundles are paired across the anterior-posterior midline of the ANC. The α tracts are situated most aborally and are the most prominent in staining for acTUBA. The δ tracts are oral to the α tracts and hug the midline. The γ and β tracts are situated on the lateral sides of the CBT, with the γ tract positioned closest to the NP and the β tract located more aborally. These four main tracts demonstrate further sub-compartments, such as those visible in the α tract on the distal slice (Fig. 2g, h) or in the β tract on the proximal slice (Fig. 2a, b). However, we could not reliably identify these additional subdivisions across all slices. This suggests either the fibers in the tracts reorganize along the proximal-distal axis (see Fig. 3h), groups of fibers are specifically lost or gained along the length of the arm, or this staining is not a reliable method for identifying further subdivisions.

**Figure 2:**
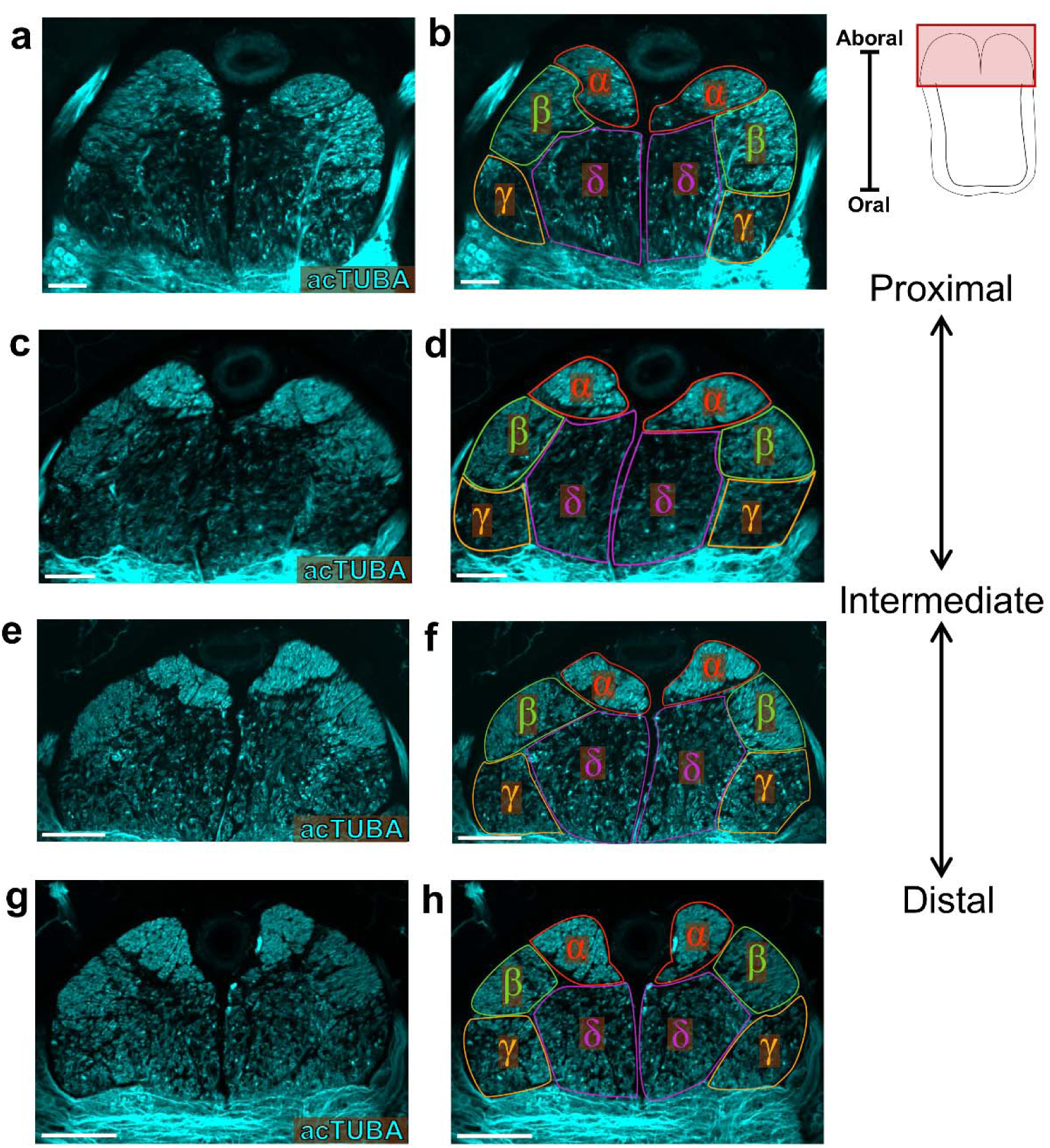
Subdivisions of the cerebrobrachial tract. **a-h,** Maximum projections of the cerebrobrachial tract (CBT) in transverse whole mount slices from proximal (**a, b**) to distal (**g, h**) stained for acetylated α-tubulin (acTUBA). The CBT can be subdivided into four major tracts which can be recognized at all proximal-distal levels: α (red), β (green), γ (orange) and δ (magenta) tracts. Data used to define the tracts is presented in the left column (**a, c, e, g**). Tracts are outlined in the right column (**b, d, f, h**). Key on top right indicates field of view. Scale bars: 100 μm. acTUBA, acetylated α-tubulin.

**Figure 3:**
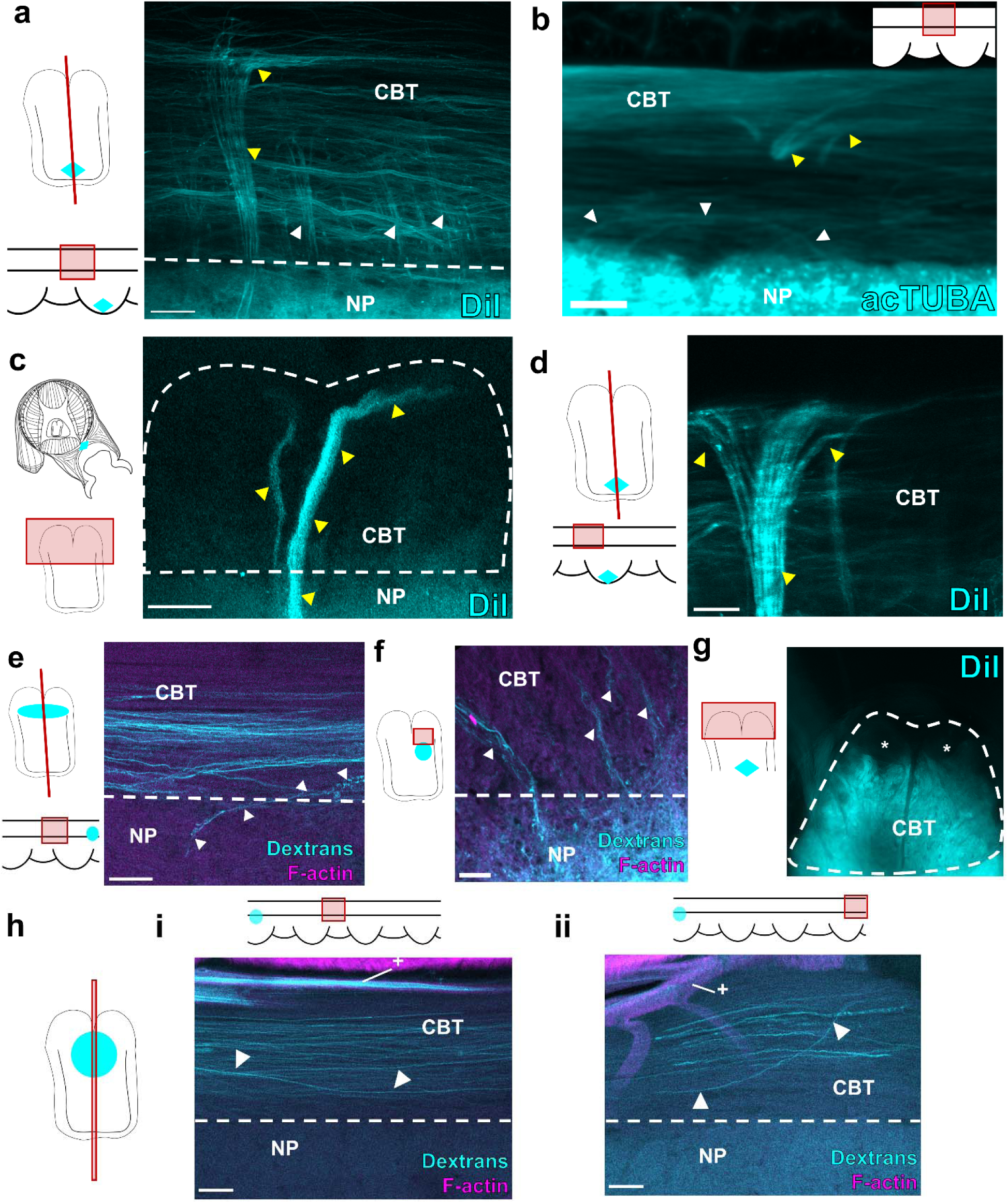
Cerebrobrachial tract and neuropil interactions. **a-b,** The cerebrobrachial tract has many interactions with the NP. Yellow arrowheads point to large fascicles. White arrowheads mark small processes. (**a**) Maximum projection of the cerebrobrachial tract (CBT) in a longitudinal wholemount with DiI (cyan) crystal placed in oral neuropil (NP). Keys indicate plane of section and field of view. Bold black lines in longitudinal key indicate the limits of the CBT. Diamond in key demonstrates location of DiI crystal. (**b**) Maximum projection of the CBT in a longitudinal wholemount immunostained with acetylated α-tubulin (acTUBA). Inset demonstrates field of view. Scale bars: 100 μm. **c,** CBT in a transverse wholemount with DiI (cyan) crystal placement in sucker musculature and including the sucker ganglion (diamond on key). Large processes that target α tract are labeled (yellow arrowheads). Scale bar: 50 μm. **d,** Maximum projection of the CBT in a longitudinal wholemount of ANC with DiI (cyan) crystal placed in oral neuropil (NP; diamond on key marks crystal placement). Connections to the α tract branch proximally and distally, indicated by yellow arrowheads. Scale bar: 100 μm. **e,** Longitudinal section of an injection of dextran (cyan) into the CBT. White arrowheads point to projection from CBT that dips into the aboral NP. Key indicates plane of section. Oval on key demonstrates injection site. Scale bar: 100 μm. **f,** Transverse section of an injection of dextran (cyan) into the aboral NP demonstrating connections to the CBT. Oval on key indicates injection site. Scale bar: 50 μm. **g,** Transverse wholemount with DiI (cyan; diamond on key) crystal placement in the aboral neuropil. The β, γ, and δ tracts are labeled, but not the α tract. **h,** Longitudinal section of an injection of dextran into the CBT. Key indicates location of injection site. Fascicles within the CBT travel a distance and rearrange. (**i**) Field of view four suckers away from the injection site. White arrowheads point to CBT fascicle that descends orally while remaining within the CBT. (**ii**) Field of view six suckers away from the injection site. White arrowheads point to CBT fascicle that ascends aborally. + denotes brachial artery. Scale bars: 100 μm. acTUBA, acetylated α-tubulin; CBL, cell body layer; CBT cerebrobrachial tract; NP, neuropil.

The CBT demonstrates many connections with the NP, with large processes targeting the aboral α tract and smaller processes targeting oral bundles (γ and δ tracts; Fig. 3a, b). By placing DiI crystals in the sucker musculature (Fig. 3c) or in the oral (sucker) territory of the ANC NP (Fig. 3d), we elicited extensive labeling of fascicles targeting to the α tract. These connections primarily targeted the CBT trunk ipsilateral to the sucker, although fibers targeting the CBT trunk contralateral to the sucker were observed (Fig. 3c). Once these processes reach the α tract, they split into proximal and distal branches, directed both towards and away from the brain (Fig. 3d). In contrast, injections of dextran into the γ and δ tracts labelled projections dipping into the aboral, brachial NP (Fig. 3e), and reciprocal injections into the aboral, brachial NP labelled projections targeting the γ and δ tracts (Fig. 3f). Lastly, DiI crystal placement into the aboral brachial NP demonstrates labeling in β, δ, and γ tracts, but not the α tract (Fig. 3g). These results suggest that the β, δ, and γ tracts correspond to connections primarily related to the brachial musculature, though connections with the sucker could also be labeled as fibers of passage. Once within the CBT, some fibers maintain their position (Fig. 3e, h). Others resort, descending orally or ascending aborally at substantial distances from the injection site (Fig. 3h). This kind of repositioning has been seen in vertebrate fiber tracts connecting the retina to the brain (Chan and Guillery, 1994).

### Longitudinal tracts in *Octopus bimaculoides* NP

We next interrogated the ANC NP for longitudinally running tracts with DiI, dextran labeling, and immunohistochemistry. The aboral, brachial NP is broadly dominated by longitudinal fibers (Fig. 4a, b). These longitudinal fibers form a prominent bilateral pair of tracts (bLTs) situated at the transition between the oral sucker NP and the aboral brachial NP (Fig. 4c). These tracts are positioned at the most aboral branch point of the oral fascicles, which feed to the oral roots innervating the sucker (Fig. 4d; Olson et al., 2025). By their location at the interface between sucker and brachial territories of the ANC, these tracts could integrate motor control information coordinating the brachial musculature and sucker movements.

**Figure 4:**
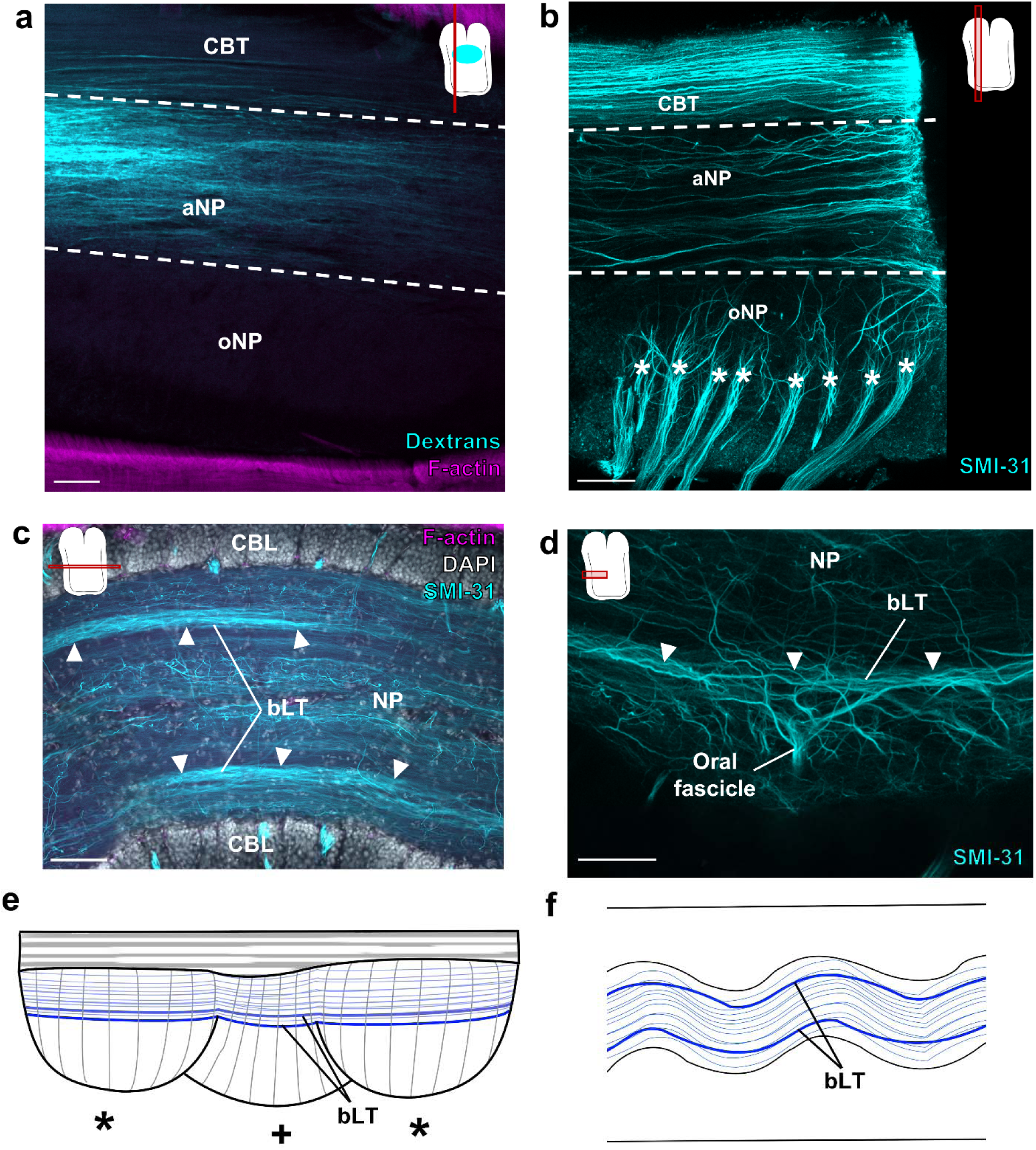
Longitudinal tracts in the aboral neuropil. **a-b,** The aboral neuropil (NP) is dominated by longitudinal fibers. **a,** Longitudinal section of an injection of dextran (cyan) into the aboral NP demonstrating longitudinal fibers. Key displays plane of section. Oval on key denotes injection site. **b,** Maximum projection of a longitudinal wholemount with SMI-31 immunostaining (cyan) marks the longitudinal fibers in the aboral NP. The oral fascicles (denoted by *) that become the oral roots targeting the sucker are also readily visible in this preparation. **c,** Horizontal section at the transition from the oral NP to the aboral NP stained for SMI-31 (cyan), F-actin (magenta), and DAPI (gray). There are two prominent bilateral longitudinal tracts (bLT), marked by white arrowheads. **d,** Maximum projection of a horizontal wholemount stained for SMI-31 (cyan) confirms that the bLT corresponds to the most aboral branch point of the oral fascicles. **e-f,** Summary diagrams showcasing the longitudinal tracts (blue) in the aboral NP in a longitudinal ANC cartoon (**e**) and a horizontal ANC cartoon (**f**). The bLT are denoted by thicker lines, and, in (**e**), are offset to demonstrate the tract on the opposite side. * in (**e**) denote ANC enlargements for suckers on the same side, + denote ANC enlargements for suckers on the opposite side. Scale bars: 100 μm. aNP, aboral neuropil; bLT, bilateral longitudinal tract; CBL, cell body layer; CBT cerebrobrachial tract; NP, neuropil; oNP, oral neuropil.

Within the oral, sucker NP, we detected an internal longitudinal tract (iLT; Fig. 5c, d) and an oral longitudinal tract (oLT; Fig. 5e, f). The trajectories of the iLT and oLT are complex, and Figure 5 a and b summarize our conclusions about their shifting relationships. In the horizontal plane, the iLT follows the internal side of the ANC, which corresponds to the internal side of the sucker (Olson et al., 2025; 2025a), switching from one edge to the other to maintain its relative positioning (Fig. 5b, d). The oLT is located oral to the iLT and sits close to the center of the oral sucker NP (Fig. 5a, e). As seen in horizontal sections, the oLT oscillates through the sucker enlargements (Fig. 5b, f). As the ANC transitions to the next sucker enlargement, the oLT bifurcates into two trunks (Fig. 5g, h). One of the oLT trunks ascends aborally to loop over the iLT, and the other continues orally (Fig. 5g, h). Both branches connect directly to the oLT in the next sucker enlargement (Fig. 5h). By projecting aborally, the oLT could receive input from other parts of the arm and mark the transition from one sucker to the next.

**Figure 5:**
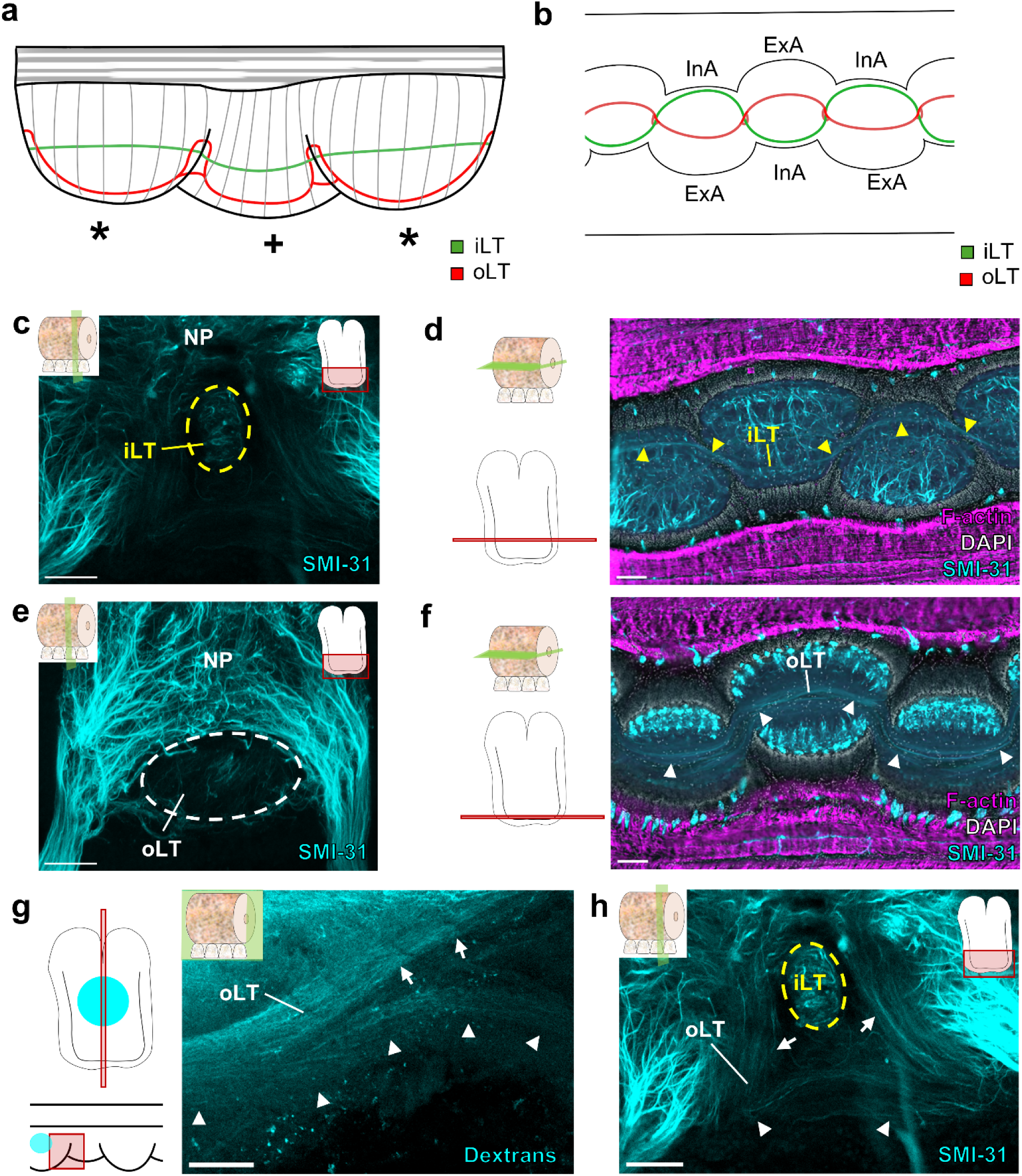
Longitudinal tracts in the oral neuropil. **a,** Schematic of a longitudinal axial nerve cord (ANC) demonstrating the trajectories of the internal longitudinal tract (iLT, green) and the oral longitudinal tract (oLT, red) as the tracts travel from sucker to sucker. * denotes ANC enlargements for suckers on the same side. + denotes ANC enlargement for suckers on the opposite side. **b,** Diagram of a horizontal section of the ANC demonstrating the paths of the oLT (red) and iLT (green) in the oral portion of the ANC. **c**, Maximum projection of a transverse ANC whole mount stained for SMI-31 (cyan). The iLT is outlined with a yellow dashed line as it crosses the midline. Key indicates transverse plane of section. Scale bar: 100 μm. **d**, Horizontal section through the oral NP stained for SMI-31 (cyan), F-actin (magenta), and DAPI (gray). The iLT, marked by yellow arrowheads, follows the internal side of the ANC. **e,** Maximum projection of a transverse ANC whole mount stained for SMI-31 (cyan). The oLT is outlined with a white dashed line. Key indicates transverse plane of section. Scale bar: 100 μm. **f,** Horizontal section through the oral NP stained for SMI-31 (cyan), F-actin (magenta), and DAPI (gray). The oLT, marked by white arrowheads, is positioned close to the midline, is located oral to the iLT, and oscillates through ANC enlargements. Key indicates horizontal plane of section. Scale bar: 150 μm. **g, h,** The oLT bifurcates at the transition between sucker enlargements. White arrowheads mark the oral branch of the oLT, and white arrows mark the aboral branch of the oLT. See loop of the oLT in (**a**). (**g**) Injection of dextran labeling the oLT at the transition between sucker enlargements. Key indicates plane of section and field of view. Oval on key indicates injection site. (**h**) Maximum projection of a transverse ANC whole mount stained for SMI-31 (cyan) at the transition between sucker enlargements. The iLT is outlined with a yellow dashed line. The aboral oLT trunk loops over the iLT. The aboral and oral oLT branches connect to the oLT on the other side. Scale bars: 100 μm. ANC, axial nerve cord; ExA, external side of the axial nerve cord; InA, internal side of the axial nerve cord; iLT, internal longitudinal tract; NP, neuropil; oLT, oral longitudinal tract.

### Comparative analysis of longitudinal tracts

We carried out a comparative analysis to assess the organization of ANC longitudinal tracts across coleoid cephalopods. We selected two species of squid for our analysis: the longfin inshore squid, *Doryteuthis pealeii*, and the hummingbird bobtail squid, *Euprymna berryi* (Fig. 6). These squid species, which diverged from octopuses 270 million years ago (mya; Albertin et al., 2022), have eight arms and two tentacles. Like the arms of *O. bimaculoides*, these arms and tentacles are muscular hydrostats (Kier and Smith, 1985; Kier, 2016). In both squid species, the tentacles are utilized in a specialized prey capture behavior: the sucker-poor tentacle stalks rapidly elongate to grab prey with the sucker-rich tentacle clubs (Hanlon and Messenger, 2018). Despite these similarities, *D. pealeii* and *E. berryi*, which diverged approximately 100 mya (Albertin et al., 2022), demonstrate key differences. *D. pealeii* are pelagic animals, found in the open ocean, whereas *E. berryi*, which are smaller animals, are benthic like *O. bimaculoides.* Additionally, the arms of *D. pealeii* have two rows of suckers (Fig. 6b), whereas the arms of *E. berryi* are lined with four rows of suckers (Fig. 6c). The tentacle clubs in *D. pealeii* are enriched with suckers similar in structure to the arm suckers, and, at the widest point, are arranged in rows of four suckers (Fig. 6b). In contrast, the clubs of *E. berryi* are covered in hundreds of minute suckers with long, tube-like stalks, and present the researchers with a Velcro-like stickiness (Fig. 6c; see also, Fig. 10a).

**Figure 6:**
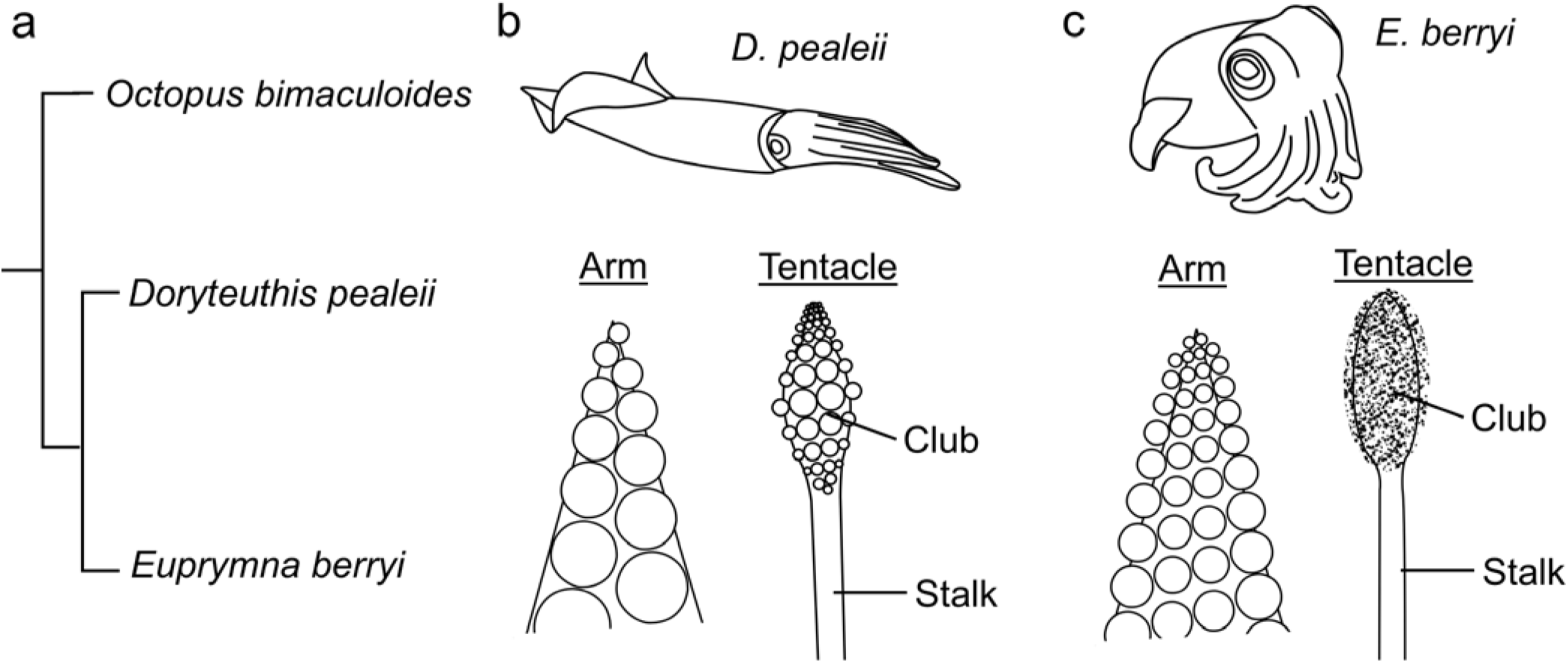
Phylogeny and gross morphology of arms and tentacles in *D. pealeii* and *E. berryi*. **a,** Tree demonstrating the relationship between *Octopus bimaculoides* and two species of squid, *Doryteuthis pealeii* and *Euprymna berryi*. Octopodiforms and decapodiforms split ∼270 million years ago (mya), *D. pealeii* and *E. berryi* split ∼100 mya. **b,** Morphology of *D. pealeii* arms and tentacle. The arm is lined with two rows of suckers, the tentacle stalk is devoid of suckers, and the tentacle club is enriched in suckers similar to the suckers on the arm. **c,** Morphology of *E. berryi* arm and tentacle. The arms are lined with 4 rows of suckers. The tentacle stalk is devoid of suckers. The tentacle club is lined with hundreds of small suckers.

#### The ANC of Doryteuthis pealeii

Like *O. bimaculoides*, *D. pealeii* has an extensive nervous system embedded in the arms and tentacles. In addition to a prominent ANC, the arms and tentacles have six intramuscular nerve cords (IMNCs) distributed around the lateral edges (Fig. 7a-c). The arms and tentacle clubs, which have suckers, also have sucker ganglia that, unlike *O. bimaculoides*, have a canonical ganglionic organization (see Olson et al., 2025a). As noted in Olson et al. (2025), the ANCs in *D*. *pealeii* do not have a clear CBT analogous to the octopus CBT, but does have major longitudinally running tracts distributed aborally and orally (Fig. 7d-f). Fibers in the aboral longitudinal tract (aLT) and the oral longitudinal tract (oLT) show a range of large diameters in transverse cross sections (Fig. 7h-k). Those in the tentacle stalk aLT are the largest (Fig. 7j). Both the aLT and the oLT are situated outside of the NP. In the arm and tentacle club, where there are suckers, the oLT oscillates around the NP (Fig. 7l, n). In the tentacle stalk, the oLT runs straight (Fig. 7m). We did not detect clear spatial subdivisions within the oLT or aLT as we did in the octopus CBT. The spatial separation of the oLT and aLT from each other hints at distinct functions. By its location closest to the suckers, the oLT may in part handle long distance sucker communication whereas the aLT may mediate long distance brachial musculature operations.

**Figure 7:**
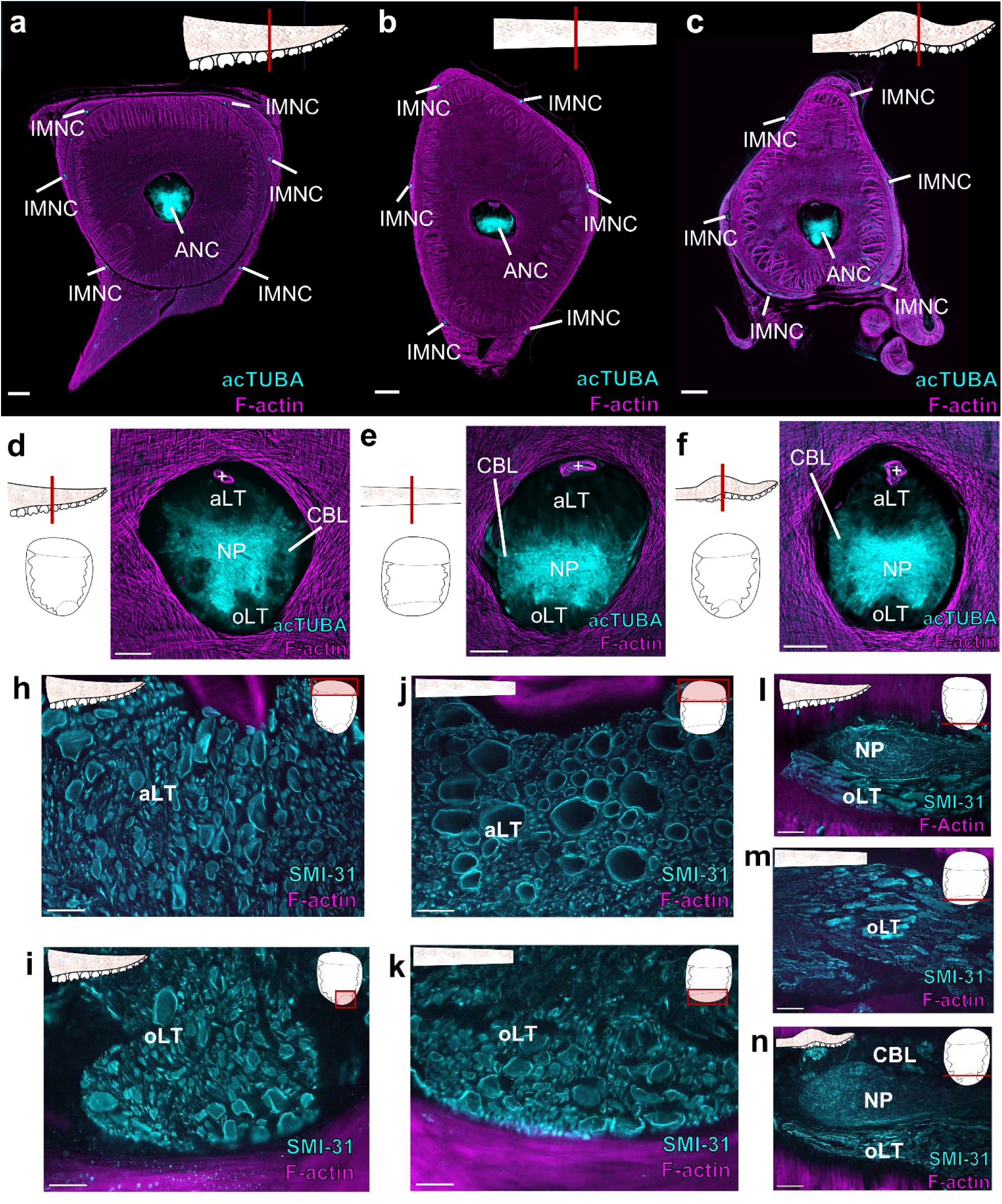
Organization of *D. Pealeii* arm and tentacle nervous system. **a-b,** Transverse section of the *D. pealeii* arm (**a**), tentacle stalk (**b**) and tentacle club (**c**) stained for acTUBA (cyan) and F-actin (magenta). The main components of the nervous system are highlighted. Scale bars: 500 μm. **d-f,** Transverse section of the axial nerve cord in the *D. pealeii* arm (**d**), tentacle stalk (**e**) and tentacle club (**f**) stained for acTUBA (cyan) and F-actin (magenta). Each ANC has an aboral longitudinal tract (aLT) and an oral longitudinal tract (oLT). + denotes brachial artery. Scale bars: 250 μm. **h-k,** The aLT and oLT are composed of large fibers. **h-i,** Transverse sections through the aLT (**h**) and oLT (**i**) in arm ANC stained for SMI-31 (cyan) and F-actin (magenta). **j-k,** Transverse sections through the aLT (**j**) and oLT (**k**) in the tentacle stalk ANC stained for SMI-31 (cyan) and F-actin (magenta). Scale bars: 50 μm. **l-n**, Horizontal sections through oLT in the arm (**l**), tentacle stalk (**m**) and tentacle club (**n**) stained for SMI-31 (cyan) and F-actin (magenta). In the arm (**l**) and tentacle club (**n**), where there are suckers, the oLT conspicuously weaves around neuropil (NP). Scale bars: 100 μm. acTUBA, acetylated α-tubulin; aLT, aboral longitudinal tract; ANC, axial nerve cord; CBL, cell body layer; INMC, intramuscular nerve cord; NP, neuropil; oLT, oral longitudinal tract.

We identified additional longitudinal fibers in the NP of *D. pealeii* ANC (Fig. 8). In the arm, these longitudinal fibers coalesce into distinct tracts situated at the interface of the cell body layer (CBL) and NP (Fig. 8a-c). The tentacle stalk ANC NP does not have distinct longitudinal NP tracts (Fig. 8d). Instead, the entire border between the CBL and NP is lined with longitudinal fibers (Fig. 8e-f). Having distinct bundles in *D. pealeii* arm but not tentacle could reflect differences in the sensorimotor demands for the suckers or the need to coordinate sucker and arm movements along a distance.

**Figure 8:**
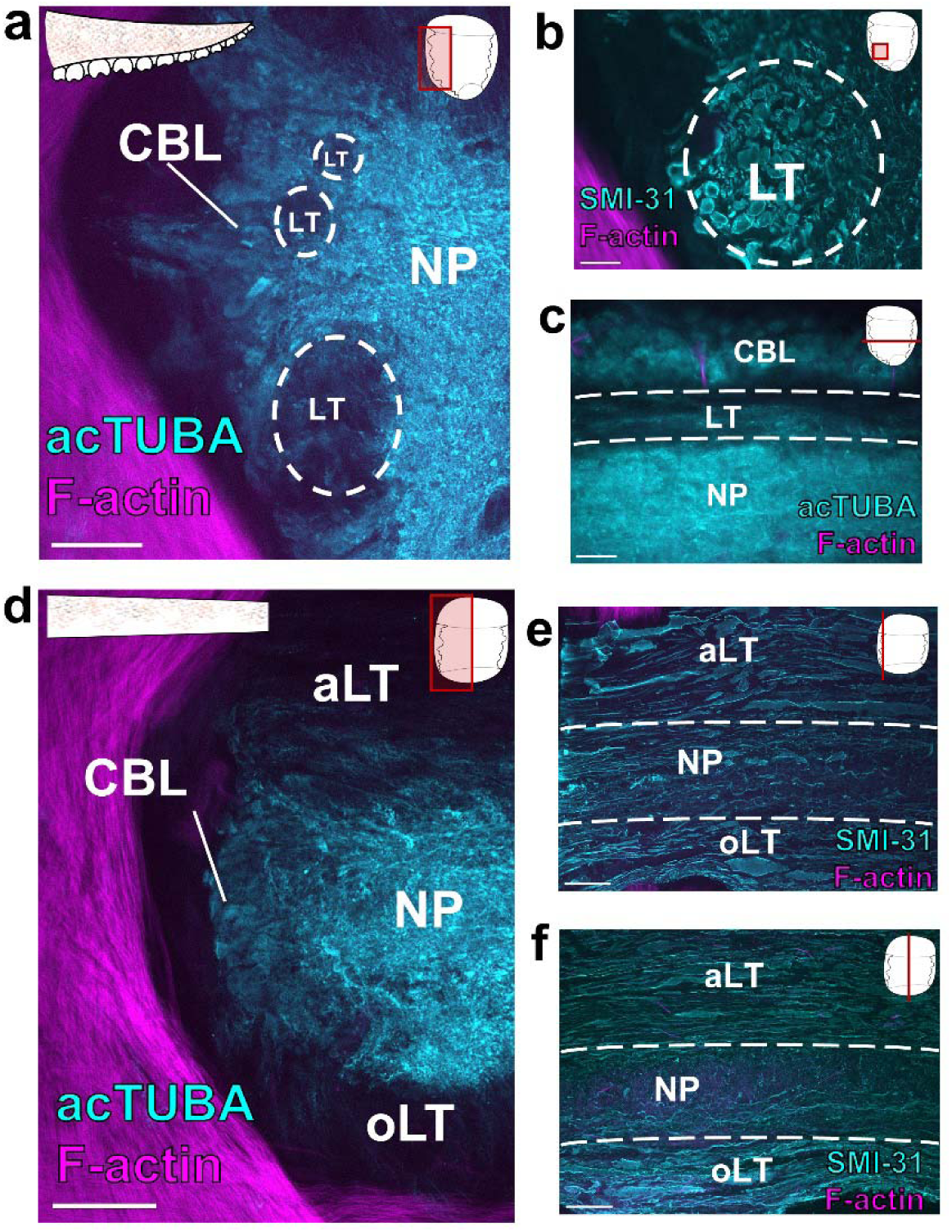
Longitudinal tracts in *D. pealeii* neuropil. **a,** Transverse section through the axial nerve cord in the squid arm stained for acetylated α-tubulin (acTUBA, cyan) and F-actin (magenta). Clear longitudinal tracts (LT), marked with white dashed line, are apparent by a decrease in acTUBA labeling. Inset depicts field of view. Scale bar: 100 μm. **b,** Transverse section through the *D. pealeii* arm ANC with SMI-31(cyan) immunostaining labels the fibers in the longitudinal tracts. Scale bar: 50 μm. **c,** Horizontal section through the *D. pealeii* arm ANC stained for acTUBA. The longitudinal tracts are located at the border of the cell body layer and neuropil. Scale bar: 50 μm. **d,** Transverse section through the ANC in the tentacle stalk stained for acetylated α-tubulin (acTUBA, cyan) and F-actin (magenta). There are no defined NP tracts. Scale bar: 100 μm. **e-f**, Longitudinal sections through the tentacle stalk ANC stained for SMI-31 (cyan) and F-actin (magenta). Longitudinal fibers are abundant at the interface of the CBL and NP (**e**) but somewhat reduce in the center (**f**). Inset displays plane of section. Scale bars: 100 μm. acTUBA, acetylated α-tubulin; aLT, aboral longitudinal tract; ANC, axial nerve cord; CBL, cell body layer; LT, longitudinal tract; NP, neuropil; oLT, oral longitudinal tract.

#### The ANC of Euprymna berryi

In the arms and tentacles of *E. berryi*, there is a prominent ANC mediating sensorimotor control (Fig. 9a-d, Fig. 10a,b). Even though the arm is lined with four rows of suckers, the ANC innervates each sucker separately, and each sucker has a dedicated sucker ganglion (Fig. 9a). *E. berryi* arms and tentacle stalks also exhibit six IMNCs (Fig. 9a, b; Fig. 10a).

**Figure 9:**
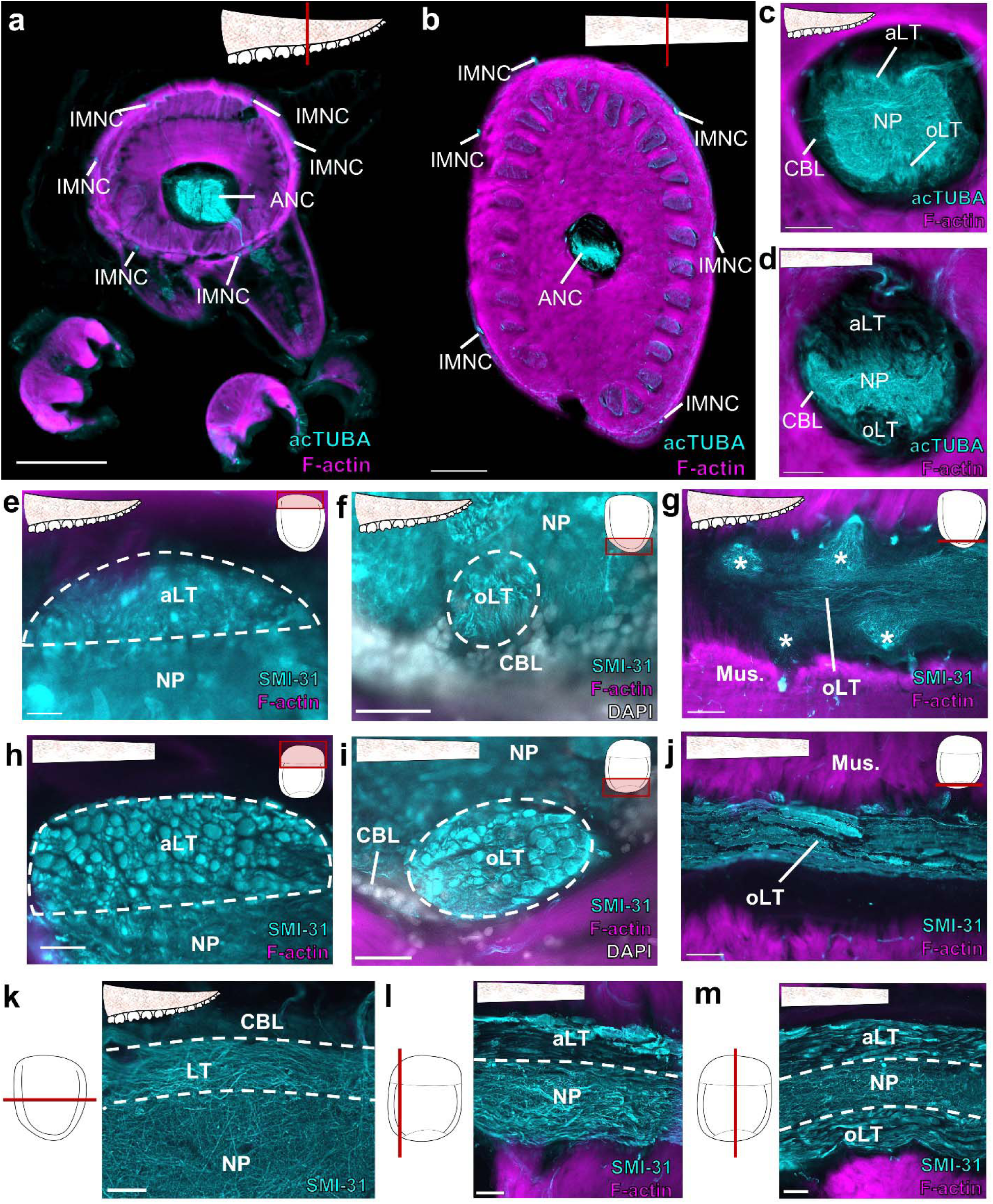
Organization of *E. berryi* arm and tentacle stalk nervous system. **a-b**, Transverse section of the *E. berryi* arm (**a**) and tentacle stalk (**b**) stained for acTUBA (cyan) and F-actin (magenta). The main components of the nervous system are highlighted. Scale bars: 500 μm. **c-d**, Transverse section of the axial nerve cord (ANC) in the *E. berryi* arm (**c**) and tentacle stalk (**d**) stained for acTUBA (cyan) and F-actin (magenta). Each ANC has an aboral longitudinal tract (aLT) and an oral longitudinal tract (oLT). Scale bars: 250 μm. **e**-**f**, The aLT and oLT in the *E. berryi* arm are composed of small fibers. Transverse sections through the aLT (**e**) and oLT (**f**) in arm ANC stained for SMI-31 (cyan), F-actin (magenta), and DAPI (gray). Notably, the arm oLT is encompassed by the cell body layer of the ANC. Scale bars: 50 μm. **g,** Horizontal section through the oLT in the arm ANC stained for SMI-31 (cyan) and F-actin (magenta). The oLT runs straight through sucker enlargements, denoted by *. Scale bar: 100 μm. **h-i,** The aLT and oLT in the tentacle stalk are composed of large fibers. Transverse section through the aLT (**h**) and the oLT (**i**) in the tentacle stalk ANC immunostained with SMI-31 (cyan), F-actin (magenta), and DAPI (gray). The oLT in the tentacle stalk is not within the cell body layer. Scale bars: 50 μm. **j,** Horizontal section through the oLT in the tentacle stalk ANC stained with SMI-31 (cyan) and F-actin (magenta). The oLT runs straight. Scale bar: 100 μm. **k-m,** Longitudinal fibers in the NP of the arm and tentacle stalk ANC are abundant at the interface of the CBL and the NP. **k,** Horizontal section through the arm ANC stained for SMI-31 (cyan). The longitudinal fibers are located at the border of the cell body layer and neuropil. Key displays plane of section. Scale bar: 50 μm. **l-m,** Longitudinal sections through the tentacle stalk ANC immunostained for SMI-31 and F-actin. Longitudinal fibers are abundant at the interface of the CBL and NP (**l**) but reduce in the center (**m**). Key displays plane of section. Scale bars: 100 μm. acTUBA, acetylated α-tubulin; aLT, aboral longitudinal tract; ANC, axial nerve cord; CBL, cell body layer; INMC, intramuscular nerve cord; NP, neuropil; oLT, oral longitudinal tract.

Within the arm of *E. berryi*, the ANC has a discernable aboral longitudinal tract (aLT; Fig. 9c, e) and oral longitudinal tract (oLT; Fig. 9c, f). The aLT is situated outside of the neuropil (Fig. 9e), whereas the oLT is within the NP (Fig. f). Both tracts are composed of small fibers. The oLT lies close to the midline and runs straight while enlargements for the suckers oscillate around it (Fig. 9g). Beyond the oLT, we failed to identify additional defined neuropil tracts. However, we found that longitudinal fibers line the interface of the CBL and the NP (Fig. 9k). With an extra-NP aboral tract and an intra-NP oral tract, the arrangement of the longitudinal tracts in the *E. berryi* arm ANC is reminiscent of those in the ANC of *O. bimaculoides*.

The tentacle stalk ANC of *E. berryi* likewise contains an aLT and oLT (Fig. 9d, h, i). Both tracts are composed of large fibers and are situated outside of the NP (Fig. 9h, i). This arrangement is similar to the distribution of longitudinal tracts in the tentacle stalk ANC in *D. pealeii*. Like in the *D. pealeii* tentacle stalk ANC, the oLT runs straight (Fig. 9j). We examined the NP of the ANC in the tentacle stalk of *E. berryi* for additional longitudinal tracts. While we failed to uncover any distinct longitudinal tracts, the border of the CBL and NP contains many longitudinal fibers (Fig. 9l-m).

Despite its unique morphology, the tentacle club in the *E. berryi* maintains major features of the cephalopod appendage nervous system: a prominent ANC and IMNCs distributed laterally (Fig. 10a). Compared with tentacle stalk, we found that the club ANC exhibits an expanded neuropil (Fig. 10b). The aLT appears compressed, and the oLT is greatly diminished (Fig. 10b,c). In major contrast to that of the tentacle stalk, the oLT is within the NP and is populated with smaller fibers (Fig. 10c). This composition of the oLT is similar to that of the *E. berryi* arm. This suggests, that within *E. berryi*, whether the ANC innervates suckers may dictate the positioning of the oLT. Indeed, the ANC in the tentacle club issues nerves that target the hundreds of minute suckers lining the club (Fig. 10d,e). The unique sensory and motor demands of this array of fine suckers warrants further study.

**Figure 10:**
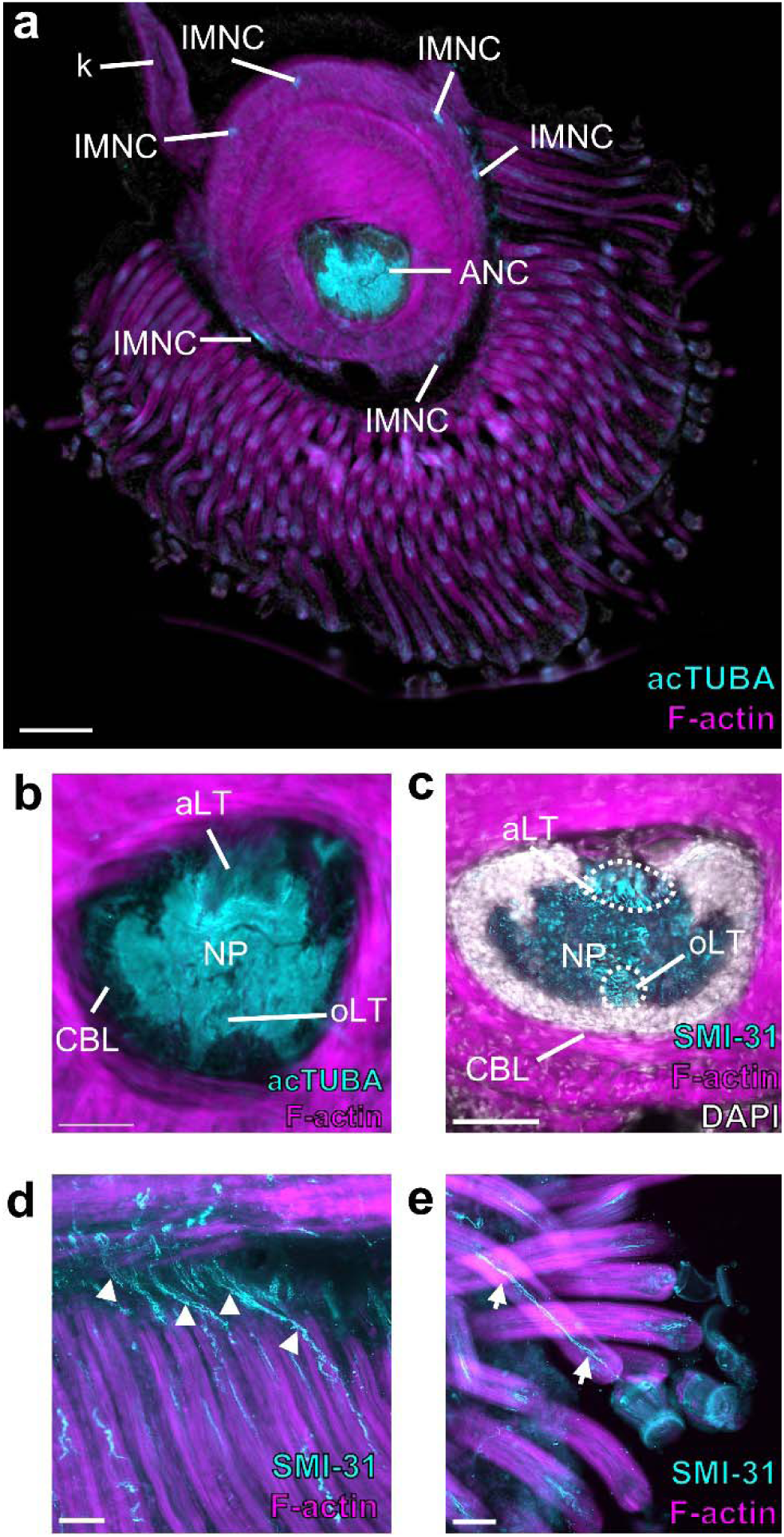
Organization of the *E. berryi* tentacle club nervous system. **a,** Transverse section of the *E. berryi* tentacle club stained for acetylated α-tubulin (acTUBA; cyan) and F-actin (magenta). The main components of the nervous system are highlighted. The tentacle club is lined with minute suckers at the end of long tube-like stalks. Scale bars: 500 μm. **b**, Transverse section of the axial nerve cord (ANC) in the *E. berryi* tentacle club stained for acTUBA (cyan) and F-actin (magenta). The neuropil (NP) is expanded and the aboral longitudinal tract (aLT) appears compressed. **c,** Transverse section of the ANC stained for SMI-31 (cyan), F-actin (magenta), and DAPI (gray). The aLT and the oral longitudinal tract (oLT) are composed of small fibers, and the oLT is within the NP. Scale bars: 250 μm. **d, e,** The ANC issues nerves that travel down the long tube-like sucker stalks to target the minute suckers. (**d**) Longitudinal section through the tentacle club stained for SMI-31 (cyan) and F-actin (magenta). White arrowheads point to ANC nerves as they begin to travel along the sucker stalks. (**e**) Longitudinal section through the tentacle club stained for SMI-31 (cyan) and F-actin (magenta). White arrows indicate an ANC nerve traveling along the sucker stalk to target the sucker. Scale bars: 25 μm. acTUBA, acetylated α-tubulin; aLT, aboral longitudinal tract; ANC, axial nerve cord; CBL, cell body layer; INMC, intramuscular nerve cord; k, swimming keel; NP, neuropil; oLT, oral longitudinal tract.

## Discussion

Many of the complex behaviors shown by the octopus require the coordination of suckers and arm musculature along the length of the arm (Kier and Smith, 1985; Gutfreund et al., 1998; Sumbre et al., 2006; Kier, 2016; Hanlon and Messenger, 2018; Kennedy et al., 2020; Bidel et al., 2022). Accordingly, the ANC in *O*. *bimaculoides* hosts many longitudinal fiber tracts that are strong candidates to support these behaviors. Some of these behaviors involve coordination of suckers along the proximal-distal axis, such as passing food along the arm according to its valence (Altman, 1971). The oLT and the α tract in the CBT may serve as a substrate for sensorimotor communication underlying these behaviors. Sucker movements often recruit the movement of the rest of the arm, whether it is to extend or rotate an arm to grasp an object or to use the arm to push an object away (Sumbre et al., 2006; Kennedy et al., 2020). The bilateral tracts situated in the brachial NP, just aboral to the sucker NP, may serve as pathways for coordinating these behaviors. It is undoubtedly the case that the different arms, together with their suckers, can be coordinated, in behaviors such as walking on the ocean floor or rotating a ball (Bennice et al., 2025; see supplemental video 1 in Olson et al., 2025). At least some of this is likely mediated by ANC-to-ANC connections traveling across arms by way of the interbrachial commissure (Graziadei, 1971; Young, 1971; Chang and Hale, 2023). Classic studies suggest that the interbrachial commissure is formed in part by longitudinal fibers from the NP of the ANC (Graziadei, 1971). Whether these longitudinal fibers arise from the NP tracts we define here and how the longitudinal tracts in the NP interconnect with the interbrachial commissure and the brain requires further study.

We confirmed the existence of the longitudinal tracts in the octopus arm nervous system with four distinct techniques— two tracing methods and two antibodies. Our results demonstrate that two features drive the organization of the longitudinal tracts: whether they are intra-neuropil tracts, such as the iLT, or extra-neuropil tracts, like the CBT, and whether they pertain to the sucker or to the brachial musculature. Within the CBT, the α tract appears to be strongly coupled with the sucker, whereas the γ and δ tracts have connections particularly to the brachial territory of the ANC. The four defined intra-NP tracts (oLT, iLT, bLTs) are either situated within the sucker territory of the ANC or demonstrate clear links to the oral roots that innervate the sucker. Although the brachial territory of the ANC contains longitudinal fibers, it lacks defined tracts with clear connections to the brachial musculature. Given its proximity to CBT, the defined tracts within the CBT, such as the γ and δ tracts could be used as intermediate relays for the brachial musculature. Nerves arising from the brachial territory of the ANC also connect to the skin covering the arm (Olson et al., 2025). Prior research suggest that efferent fibers from the chromatophore lobes in the brain directly connect to the arm chromatophores (Young, 1971). These fibers most certainly travel through the CBT before descending into the brachial NP to join the nerves innervating the skin. Additional studies of how the subtracts of the CBT interconnect with the brain would reveal the pathways used by chromatophores efferent fibers.

In the vertebrate nervous system, spinal cord tracts are divided into sensory tracts and motor tracts. Previous work in the octopus arm nervous system demonstrates that broad segregation of sensory and motor neuron cell bodies is not a principle feature of the ANC CBL and suggests that ANC nerves are mixed, carrying both sensory and motor information (Rossi and Graziadei, 1958; Rowell, 1966; Gutfreund et al., 2006; Olson et al., 2025; 2025a). The longitudinal tracts within the ANC could be subdivided into separate bundles for sensory and motor information, providing a method for unmixing sensory and motor signals. Functional studies are needed to assess this possibility. Physiological studies have shown that mechanosensory signals, at the least, travel both proximally and distally along the ANC (Gutfreund et al., 2006; Chang and Hale, 2023). This aligns with our tract-tracing results, which demonstrate proximal and distal connections to the longitudinal tracts.

A comparison to squid demonstrates that despite differences in appendage morphology, environmental context and behavioral needs, there are shared features in ANC longitudinal tract structure. Across the cephalopod species studied, all ANCs contain an aboral longitudinal tract (aLT; CBT in *O. bimaculoides*) located outside of the NP and an oral longitudinal tract (oLT). With its two trunks and readily observable subdivisions, the CBT in *O. bimaculoides* demonstrates a more complex organization when compared with the aLT in squid, which could underlie an increase in behavioral complexity and sensory function (Albertin et al., 2022; Kang et al., 2023). The oLT in the *D. pealeii* arm is composed of large fibers and is situated outside of the NP, whereas the oLT in the *O. bimaculoides* and *E. berryi* arms is solidly within NP. This displacement of the oLT in *D. pealeii* suggests a bias towards longer distance connections. Whether oLT or aLT in squid correspond to the CBT by interconnecting with the brain needs experimental testing with tract-tracing methods.

The organization of the ANC within the tentacle stalk appears remarkably conserved between *D. pealeii* and *E. berryi*. In each, the oLT and aLT are extra-NP tracts, composed of fibers with large diameters. Additional longitudinal fibers dominate the interface between the CBL and NP. This similarity in neural structure is underscored by similarities in the behavioral output in tentacle stalks (Kier and Smith, 1985; Kier, 2016). The tentacle stalks elongate rapidly to grab prey, reaching the target in 15-35 ms in loliginid squid (Kier, 1985). With the speed of this motor act, the main requirement of the tentacle stalk ANC could simply be to propagate a signal from the brain along the length of the stalk and to relay sensory feedback from the clubs to the brain. In the absence of myelin, large nerve fibers in the aLT and oLT within the stalk ANC seem well suited for fast transmission of a neural signal.

The longitudinal tracts in the ANCs of octopus and squid arms exist in the context of a clear segmental organization of the ANC cell body layer (Olson et al., 2025). Neither the intra-NP nor extra-NP tracts themselves are segmented, but, by their positioning, the intra-NP tracts would likely receive modulation from the neurons situated in the segments. The CBT, as an extra-NP tract, may serve as an unmodulated relay of sensorimotor commands. Further descriptions of the anatomical and functional inputs to the intra-NP and extra-NP tracts would clarify how neural signals are transformed along the tracts.

## Conclusion

Our findings point to a hierarchy within the arm nervous system for sensorimotor control of the suckers and the brachial musculature. First, each sucker houses a sucker ganglion, a neural center well situated for local sensorimotor integration for the sucker with relays to the oral ANC (Olson et al., 2025a). The ANC contains enlargements for each sucker, which are subdivided into segments to create a topographic map, or “suckerotopy”, for each sucker (Olson et al., 2025). This establishes a further level of sensorimotor control. Next, as described in this study, multiple complex longitudinal circuits within the NP interconnect each ANC enlargement to the next, allowing for coordinated movements across the suckers and brachial musculature. Finally, there is the CBT, the massive octopus aboral fiber tract, which provides for longer distance intra-arm connections for both the sucker and the brachial neuropil. This hierarchy may be key for understanding how the interbrachial connectives and intermediate motor areas within the brain mediate sensorimotor control for octopus arm movements, from simple deflections of individual suckers to complex, coordinated whole body actions.

## Acknowledgements

We thank Dr. Chuck Winkler of Aquatic Research Consultants for providing us with octopuses, and Dr. Caroline Albertin, Dr. Thea Rodgers and Ms. Natalie Grace Schulz for providing us with squids. We extend special thanks to Ms. Aashna Moorjani for invaluable assistance with analysis of octopus data and the processing *Euprymna berryi* samples and for comments. We thank Ms. Amelia Cheng, Mr. Jan Kasal, and Ms. Aashna Moorjani for assistance with tracer injections and tissue processing. Imaging was performed at the University of Chicago Integrated Light Microscopy Core (RRID: SCR_019197). We thank Dr. Christine Labno and Mr. Khalil Rodriguez for their invaluable assistance. This work was supported by the NIH UF1NS115817 award (CWR).

## Data availability

Data available upon request

